# Possibilities for improvement of humoral innate immunity in turkeys and hens in conditions of thermal stress by the immunomodulator Immunobeta

**DOI:** 10.1101/305300

**Authors:** N. Bozakova, L. Sotirov, S. Denev, Ts. Koynarski

**Author notes:** Corresponding author at: Dept. Animal Genetics, Faculty of Veterinary Medicine, Trakia University, 6000 Stara zagora, Bulgaria. E-mail address (L.Sotirov).

## Abstract

The experiments for evaluation of the effect of Immunobeta were performed with turkeys and organically reared layer hens. Turkeys received Immunobeta at a dose of 4 g/kg feed during the beginning of lay (cold period), thermoneutral period and hot summer period. In hens, two dietary levels of Immunobeta were tested: 2 g/kg and 4 g/kg. Blood serum and egg lysozyme levels, alternative pathway of complement activation (APCA) and beta-lysins concentrations were determined. It was found out that Immunobeta had a beneficial effect on serum lysozyme, APCA activity and beta-lysins in turkeys and hens. The lysozyme concentration in egg white was statistically significantly higher in eggs produced by birds treated with both doses of the immunomodulator.

## Introduction

In modern society, the welfare aspects of birds’ prophylaxis and treatment become especially important. During the last two decade, there are a number of potential non-therapeutic alternatives of antibiotic growth promoters (AGPs) in animal nutrition and production of safety (antibiotic-free) products, including probiotics, prebiotics, symbiotics, immunostimulants, direct feed microbials and etc. (Denev, 1996, 2006, 2006a, 2008; Denev *et al*., 2000, 2006b, 2009; Staykov *et al*., 2007; Huyghebaert *et al.*, 2011). A lot of them demonstrated positive results, equivalent to the AGPs, but without the stigma of increasing antimicrobial-resistant bacteria. They stimulated probiotic intestinal flora, ecological balance, inhibition of pathogens, humoral immunity, gut health, animal welfare, growth performance, quality and safety of animal products (Denev, 2008; Zhang and Kim, 2014; Chen *et al.*, 2017).

There are no reports about the effects of the immunomodulator Immunobeta on the innate immunity of turkeys and layer hens at the time of beginning of lay as well as in birds under heat stress. The purpose of these experiments were to investigate the effects of the immunomodulator Immunobeta on serum lysozyme concentrations, alternative pathway of complement activation and beta-lysins in turkeys and organically reared layers in conditions of environmental thermal stress.

## Material and Methods

### Experimental design

The effects of Immunobeta were evaluated on turkeys and hens in a free-range system during 2015-2016. The birds were treated with Immunobeta as per recommendations of the manufacturer Chemifarma, Italy. The immunostimulatory preparation was produced by enzymatic autolysis of selected yeast strains (*Saccharomyces cerevisiae*) followed by natural extraction of components of yeast cells. The immunomodulator contains three important components: beta-glucans (30%), mannanoligosaccharides (25%) and nucleotides (5%). The experiments for evaluation of immunomodulator’s effects in turkey breeders and organic hens were performed during different periods of the year: beginning of lay (cold period), thermoneutral period and hot summer period.

1. The study on the effect of the immunomodulator on turkey breeders took place in the turkey farm of the Agricultural Institute, Stara Zagora with 30 turkey breeders (15 experimental and 15 control birds). The influence of the preparation was tested from February to August 2015. Tested birds received the immunomodulator Immunobeta at a dose of 4.0 g/kg feed during the beginning of egg lay (February-March 2015, cold period), the thermoneutral period (April-May 2015) and the hot summer period (June-July 2015). During these periods, the blood serum and egg white lysozyme, alternative pathway of complement activation (APCA) and beta-lysins were determined. The duration of the experiment was 7 months.
2. The study on the effect of Immunobeta on humoral immunity of hens reared in a free-range system was performed in the Experimental farm of the Agrarian University — Plovdiv from November 2015 (19 weeks of age) to July 2016 (54 weeks of age). The tests were carried out on 75 dual-purpose rural hens Tetra Super Harco (originating in Babolna Tetra Kft, Hungary), housed in sleeping houses and walking yards. The effects of two dietary Immunobeta levels were followed out: 2.0 g/kg and 4.0 g/kg. The birds were divided in three groups: Group I (control), Group II (supplemented with 2.0 g/kg) and Group III (supplemented with 4.0 g/kg). The effect of the immunomodulator was monitored during three subperiods: beginning of egg lay (January-March 2016, cold period), thermoneutral period (April-May 2016) and hot summer period (June-July 2016). During these periods, the blood serum and egg white lysozyme, alternative pathway of complement activation (APCA) and beta-lysins were determined.

### Assay methods

Serum lysozyme concentrations were determined by method of Lie (1985), alternative pathway of complement activation (APCA) by method of Sotirov (1986) and beta lysins by method of Buharin *et al*. (1977).

### Statistical analysis

Data were statistically processed by on-way analysis of variance (ANOVA) using statistical software GraphPad InStat 3.06 at a level of significance P<0.05.

## Results

The results about the effect of Immunobeta supplementation on serum lysozyme, APCA activity and beta-lysins in turkey breeders are presented in Table 1. Under the influence of Immunobeta lysozyme activity tended to increase during all three subperiods. This factor of innate immunity was increased statistically significantly during the thermoneutral period (P< 0,001). A similar tendency occurred in the time course of APCA after supplementation with the immunomodulator. Although not statistically significant, the differences between control and experimental birds marked a tendency for higher serum APCA in supplemented birds at the time of beginning of lay. The beta-lysins percentage also tended to be higher during the beginning of lay and during the thermoneutral period despite the insignificant differences.

**Table 1.** Effect of Immunobeta on serum lysozyme concentrations, APCA and beta lysines in

| Periods | Lysozyme (mg/L) |  | APCA (CH50) |  | Beta lysines (%) |  |
| --- | --- | --- | --- | --- | --- | --- |
|  | Controls | Immunobeta<br>4g/kg | Controls | Immunobeta<br>4g/kg | Controls | Immunobeta<br>4g/kg |
| Cold | 0,96±0,29 | 1, 13±0,27 | 692,78±24,54 | 695,90±9,84 | 34,7± 2,9 | 47,8 ± 5,5 |
| Termoneutral | 0,32±0,05 | 0,72±0,11** | 619,45±29,34 | 632,54±24,83 | 47,6± 5,01 | 57,9± 6,8 |
| Hot | 0,31±0,03 | 0,35±0,04 | 468,55±11,06 | 488,66±9,36 | 56,9±1,6 | 56,5±1,9 |

turkey breeders.
\*\* - P<0.01

As egg white lysozyme was concerned, there was no positive effect of Immunobeta on this parameter in experimental turkeys (Table 2).

**Table 2.** Effect of Immunobeta on egg white lysozyme concentrations in turkey breeders

| Periods | Controls | Immunobeta 4g/kg | (mg/L). |
| --- | --- | --- | --- |
| Cold | 499,33±37,19 | 511, 04±17,98 |  |
| Termoneutral | 532,67±59,55 | 475,09 ± 10,01 |  |
| Hot | 587,83±51,83 | 306,51±21,59** |  |

\*\* - P<0.01

The results summarizing the effects of Immunobeta on serum lysozyme, APCA and beta-lysins in organic layer hens are shown in Table 3. Lysozyme concentrations decreased parallelly to increase in Immunobeta dose and during the hot period, the reduction is statistically significant (P<0.05). This dose-dependent change confirmed the effects of Immunobeta on the innate humoral immunity of hens.

**Table 3.** Effect of Immunobeta on serum lysozyme concentrations, APCA and beta lysines in hens.

| Groups | Lysozyme<br>(mg/L) | APCA<br>(CH50) | Beta lysines<br>(%) |
| --- | --- | --- | --- |
| Cold period |  |  |  |
| Controls | 1,71±0,22 | 543,77±28,15 | 40,56±1,29 |
| 0,2% Immunobeta | 1,50±0,25 | 567,12±25,96 | 48,99±5,62 |
| 0,4% Immunobeta | 1,53±0,15 | 613,74±26,39* | 53,17±3,23** |
| Termoneutral period |  |  |  |
| Controls | 1,88±0,35 | 493,77±32,67 | 40,56±2,22 |
| 0,2% Immunobeta | 1,50±0,41 | 533,79±24,84 | 52,32±5,82* |
| 0,4% Immunobeta | 1,53±0,49 | 563,74±14,48* | 50,34±3,90* |
| Hot period |  |  |  |
| Controls | 2,05±0,50 | 443,77±18,99 | 38,90±4,35 |
| 0,2% Immunobeta | 1,33±0,39 | 464,45±31,19 | 30,65±3,51 |
| 0,4% Immunobeta | 0,87±0,29* | 530,41±10,13*# | 33,67±4,01 |

\*- P<0.05; \*\*- P<0.01; #- statistical difference between group treated with 0,4% *Immunobeta* and control group in the hot period.

A relevant proof for the improved resistance of free-range hens was the change in APCA activity in blood. There was a statistically significant increase in APCA activity during the three subperiods after supplementation with 4.0 g/kg immunomodulator (P<0.05). A similar trend for higher APCA activity was also observed in hens treated with 2.0 g/kg Immunobeta as a dietary supplement although the differences vs controls were not significant.

Table 4 presents the results for egg white lysozyme concentrations in organic layer hens under the influence of Immunobeta. They were statistically significantly higher in eggs produced by hens treated with 4.0 g/kg Immunobeta (P<0,05). A similar change was noted in eggs produced by hens treated at 2.0 g/kg immunomodulator - the differences vs controls was statistically significant during the hot summer period (P<0,05).

**Table 4.**
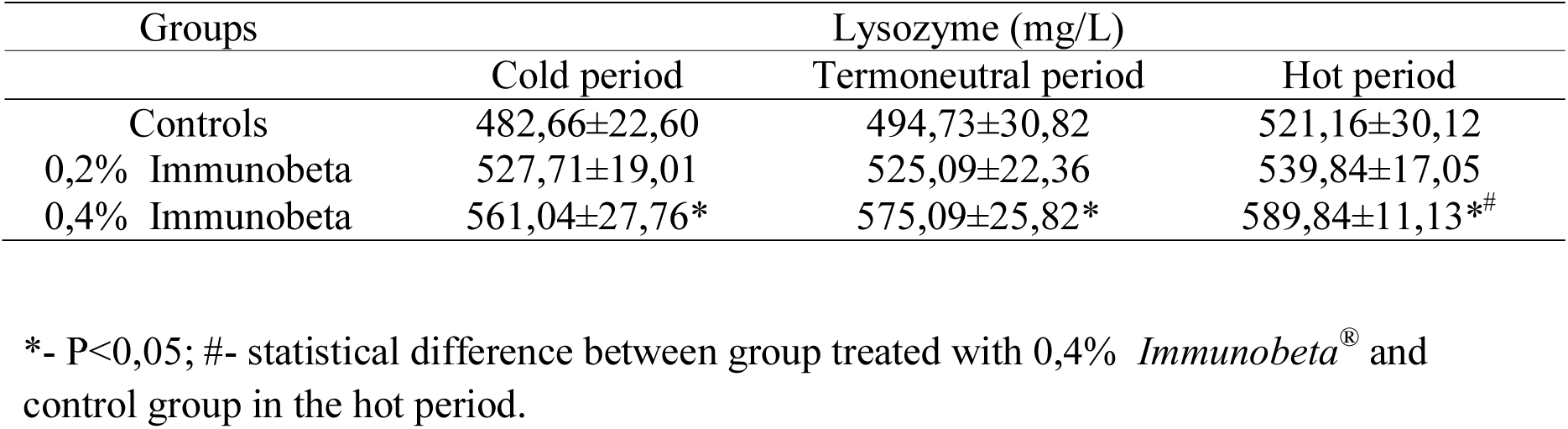
Effect of Immunobeta on egg white lysozyme concentrations in hens.

## Discussion

The results about the effect of Immunobeta on parameters of natural humoral immunity in studied birds affirmed categorically the positive effect of the immunomodulator *Immunobeta*^®^. Iimprovements of gastrointestinal morphology, physiology, microbiology and growth performance are reported by Denev (2006b, 2008), Yin *et al*. (2008), Al-Mansour *et al*. (2011), Silva *et al*. (2009), Fathi *et al*. (2012) and Bozakova *et al*. (2016) in broiler chickens, and by Huff *et al*. (2013) in turkeys. Other studies have evidence enhanced humoral immunity, resistance to diseases and lower death rates in birds after dietary supplementation of yeasts (Gao *et al.*, 2008; Yin *et al.*, 2008; Fathi *et al.*, 2012). Bozakova *et al*. (2012) demonstrated that the welfare of chicken breeders during the hot summer period has been substantially improved through supplementation of zinc (100 mg/kg Zinteral 35, Lohmann Animal Health, Germany), containing 35 mg zinc/kg as zinc oxide) and zinc + 250 mg/kg vitamin C (100 mg/kg Zinteral 35 together with 250 mg/kg vitamin C). Bozakova *et al*. (2013a) confirmed the beneficial effects of these dietary supplements on the welfare of New Hampshire breeders reared during the cold winter period. Similar positive influence was reported after supplementation of 1% L-arginine, Zinteral 35, 250 mg/kg vitamin C and 1% Arginine and 250 mg/kg vitamin C to the feed of turkeys and chicken breeders both during the hot and cold periods of the year (Bozakova *et al.*, 2009; Bozakova *et al.*, 2013b; Bozakova *et al*., 2014). Sotirov *et al*. (2000) and Denev (2008) observed a positive effects of the probiotic Lacto-Sacc^®^ and at a lesser extent of lactose (as prebiotic) on the serum lysozyme concentrations and the alternative pathway of complement activation in broiler chickens. In turkeys treated with the above feed supplements, Sotirov *et al*. (2001) obtained similar results for the same factors of non-specific resistance. Sotirov *et al*. (2007) established also the stimulating effect of organic selenium (Sel-Plex^®^) in sows and their progeny, on blood serum lysozyme concentrations and complement levels. Karakolev *et al*. (2013a,b) and Gospodinova *et al*. (2013) provided proofs for significant stimulating effect of the preparation Helpankar on serum lysozyme, gamma interferon and alternative pathway of complement activation in layer hens. Zhang *et al*. (2012) established that a dietary supplement containing yeast cellular wall products boosted the immune system of broiler chickens treated with the immunosuppressing agent cyclosporine A. Furthermore, Sadeghı *et al*. (2013) reported about a positive effect of a dietary prebiotic based on mannan-oligosaccharides and beta-glucans on the immune response of infected chickens. Comparable data about favourable influence of mannan-oligosaccharides and beta-glucans on immune performance of chickens are also reported by Shao *et al*. (2013), Shanmugasundaram *et al*. (2013), Huff *et al*. (2013). Similar data have been published by Valchev *et al*. (2015) in ducks. According to Czech *et al*. (2014) the dietary supplementation with 6% *Yarrowia lipolytica* yeast increased lysozyme levels in turkey breeders. Results about increased egg white lysozyme concentrations are reported in Lohmann Brown hens, treated with 0.4% (4.0 g/kg feed) Immunobeta in previous studies of ours (Bozakova *et al.*, 2017). Bozakova *et al*. (2016) reported that immunomodulator Immunobeta, applied as dietary supplement at doses of 3.0 and 4.0 g/kg feed, stimulated the intestinal villi height and outer diameter of glandular crypts in the small intestine of broiler chickens. The application of the preparation at doses of 2.0 and 4.0 g/kg had beneficial effects on the growth of epithelium lining the glandular crypts and adjacent intestinal villi.

## Conclusions

Immunobeta, offered at a dose of 4.0 g/kg feed had a beneficial effect on natural humoral immunity factors - it increased significantly lysozyme levels in turkeys, enhanced APCA activity and beta-lysins in hens. Applied at a dose of 4.0 g/kg, the tested immunomodulator increased significantly egg white lysozyme concentrations in hens, presuming a higher shelf life of eggs and possibly, better protection of chick embryos.

## References

Al-Mansour, S., A. Al-Khalf, I. Al-Homidan, and M.M. Fathi (2011). Feed efficiency and blood hematology of broiler chicks given a diet supplemented with yeast culture. International Journal of Poultry Science, 10, 603–607.

Bozakova, N.A., D. Girginov, and A. Atanasov (2013a). Welfare of Barred Plymouth Rock poultry flocks supplemented with zinc, vitamin C and L-arginine during the hot summer period using a mathematical assessment model. Proceeding of the 16-th ISAH Congress, 5-9 May, 2013, Nanjing, China, pp. 23–26.

Bozakova, N.A., M. Oblakova, K. Stoyanchev, I. Yotova, and M. Lalev (2009). Ethological aspects of improving the welfare of turkey breeders in the hot summer period by dietary L-arginine supplementation. Bulgarian Journal of Veterinary Medicine, 12, 3, 185–191.

Bozakova, N.A., D. Dimitrov, L. Sotirov, P. Petrov, V. Gerzilov, and Ts. Koynarski (2016). Effect of immunomodulator Immunobeta on histological features of intestinal villi and crypts in broiler chickens. Ciencia e Tecnica Vitivinicola, 31, 4, 141–149.

Bozakova, N.A., S. Popova-Ralcheva, V. Sredkova, V. Gerzilov, A. Atanasov, and N. Georgieva (2012). Mathematical welfare assessment model of chicken breeder flocks. Bulgarian Journal of Agricultural Science, 18, 2, 278–287.

Bozakova, N.A., V. Gerzilov, A. Atanasov, and I. Chukacheva (2013b). Welfare assessment of breeder hens supplemented with zinc and vitamin during the cold winter period. Bulgarian Journal of Veterinary Medicine, 16, 3, 170–178.

Bozakova, N.A, L., Sotirov, Ts. Koynarski, and D. Gundasheva (2017). Effect of immunomodulator Immunobeta on humoral innate and acquired immune response in layer hens. Pakistan Veterinary Journal (In Press).

Bozakova, N.A, and V. Gerzilov (2014). Opportunities for the Welfare Improvement of Laying Hens under Semi-open Rearing during the Cold Period with Arginine and Vitamin C Supplementation. Turkish Journal of Agricultural and Natural Sciences, Special Issue, 1, 793–798.

Buharin, O.V., A.P. Luda, and R.I. Bigeev (1977). Fotonefelometricheskij metod opredelenia beta-lizinov v sivarotke krovi. In: Sistema beta-lizina I ee rol v klinicheskoj I eksperimentalnoj medicine (Karpov S.P., ed): Izdatelstvo Tomskogo Universiteta, Tomsk, USSR, pp 52–53.

Czech, A., M. Merska, and K. Ognik (2014). Blood immunological and biochemical indicators in turkey hens fed diets with a different content of the yeast Yarrowia lipolytica. Ann. Anim. Sci., 14, 4, 935–946.

Chen, C. Y., S.W. Chen, and H.T. Wang (2017). Effect of supplementation of yeast with bacteriocin and *Lactobacillus* culture on growth performance, cecal fermentation, microbiota composition, and blood characteristics in broiler chickens. Asian-Australas Journal of Animal Science, 30, 2, 211–220.

Denev, S.A., Y. Staykov, R. Moutafchieva, and R.G. Beev (2009). Microbial ecology of the gastrointestinal tract of fish and the potential application of probiotics and prebiotics in finfish aquaculture. International Aquatic Research, 1, 1–29.

Denev, S.A. (2008). Ecological alternatives of antibiotic growth promoters in the animal husbandry and aquaculture. D.Sc. Thesis, Department of Biochemistry and Microbiology, Trakia University, Stara Zagora, Bulgaria, pp. 294.

Denev, S.A. (2006). Role of Lactobacilli in Gastrointestinal Ecosystem. Bulgarian Journal of Agricultural Science, 12, 1, 63–114.

Denev, S.A. (2006a). Effect of different growth promoters on the cecal microflora and performance of broiler chickens. Bulgarian Journal of Agricultural Science, 12, 3, 461–474.

Denev, S.A., I. Dinev, I. Nikiforov, and V. Koinarski (2006b). Effect of Mananoligosaccharides on Composition of the Cecal Microflora and Performance of Broiler Chickens. Bulgarian Journal of Ecological Science, 5, 1, 10–16.

Denev, S.A., I. Suzuki, and H. Kimoto (2000). Role of Lactobacilli in Human and Animal Health. Animal Science Journal, 71, 6, 549–562.

Denev, S.A. (1996). Probiotics-Past, Present and Future. Bulgarian Journal of Agricultural Science, 2, 445–474.

Fathi, M. M., S. Al-Mansour, A. Al-Homidan, A. Al-Khalaf, and M. Al-Damegh (2012). Effect of yeast culture supplementation on carcass yield and humoral immune response of broiler chicks. Veterinary World, 5, 11, 651–657.

Gao, J., H.J. Zhang, S.H. Yu, S.G. Wu, I. Yoon, J. Quigley, Y.P. Gao, and G.H. Qi (2008). Effects of yeast culture in broiler diets on performance and immunomodulatory functions. Poultry Science, 87, 1377–1384.

Gospodinova, K., K. Rumen, L. Sotirov, M. Bonovska, and A. Angelov (2013). Quantitative assessment of Interferon in blood serum of layer hens following treatment with polybacterial immunomodulator. 8th Balkan Congress of Microbiology “Microbiologia Balkanika’ 2013”, October 2-5, 2013, Veliko Tarnovo, Abstract VM13, pp. 89.

Huyghebaerta, G., and R.F. Ducatelle van Immerseel (2011). An update on alternatives to antimicrobial growth promoters for broilers. Veterinary Journal, 187, 182–188.

Huff, G.R., W.E Huff, and S. Jalukar (2013). The effects of yeast feed supplementation on turkey performance and pathogen colonization in a transport stress/*Escherichia coli* challenge. Poultry Science, 92, 655–662.

Karakolev, R., L. Sotirov, and M. Bonovska (2013a). Quantitative assessment of lysozyme and complement in blood serum of layer hens treated with polybacterial immunomodulator. 8th Balkan Congress of Microbiology “Microbiologia Balkanika’ 2013”, October 2-5, 2013, Veliko Tarnovo, Bulgaria, VM14, pp. 89.

Karakolev, R., L. Sotirov, M. Bonovska, K. Gospodinova, D. Nikolov, A. Angelov, Ts. Koynarski, and P. Petkov (2013b). Influence of age, technologies of growing and polybacterial immunomodulator on serum lysozyme concentrations and complement activity in laying hens. 8th Balkan Congress of Microbiology “Microbiologia Balkanika’ 2013”, October 2-5, 2013, Veliko Tarnovo, Bulgaria, Abstract VM15, pp. 90.

Lie, O., H. Solbu, and M. Sued (1985). Improved agar plate assays of bovine lysozyme and haemolytic complement activity. In: Markers for resistance to infection in dairy cattle. Ph.D. Thesis, National Veterinary Institute, Oslo, Norway, pp 1–12.

Sadeghı, A.A., A. Mohamma, P. Shawrang, and M. Amınafshar (2013). Immune responses to dietary inclusion of prebiotic-based mannan-oligosaccharide and β-glucan in broiler chicks challenged with Salmonella enteritidis. Turkish Journal of Veterinary and Animal Sciences, 37, 206–213.

Shanmugasundaram, R., M. Sifri, and R.K. Selvaraj (2013). Effect of yeast cell product (CitriStim) supplementation on broiler performance and intestinal immune cell parameters during an experimental coccidial infection. Poultry Science, 92, 358–363.

Shao, Y., Y. Guo, and Zh. Wang (2013). β-1,3/1,6-Glucan alleviated intestinal mucosal barrier impairment of broiler chickens challenged with Salmonella enterica serovar Typhimurium. Poultry Science, 92, 1764–1773.

Silva, V.K., J. Della Torre da Silva, K.A.A. Torres, D.E. de Faria Filho, H. Hirota Hada, and V.M. Barbosa de Moraes (2009). Humoral immune response of broilers fed diets containing yeast extract and prebiotics in the prestart phase and raised at different temperatures. Journal of Applied Poultry Research, 18, 530–540.

Sotirov, L., D. Goundasheva, M. Andonova, S. Denev, M. Dimitrov, and N. Vassilev (2007). Sel-Plex^®^ and sodium selenite dietary supplements with resulting serum lysozyme and complement activities in sows and progeny during post-partum periods. Trakia Journal of Sciences, 5, 1, 20–27.

Sotirov, L., S. Denev, I. Tsachev, M. Lalev, M. Oblakova, and Z. Porfîrova (2001). Effect of different growth promoters on lysozyme and complement activity. II. Studing in turkeys. Rev. Med. Vet., 152, 1, 67–70.

Sotirov, L., S.A. Denev, And V.K. Georgieva (2000). Effect of different growth promoters on lysozyme and complement activity of broiler chicks. Bulgarian Journal of Agricultural Science, 6, 75–82.

Sotirov, L. (1986). Method for determination of the alternative pathway of complement activation in some animals and man. In: Forth Scientific Conference of Agriculture, Stara Zagora, Bulgaria, pp. 1–10.

Staykov, Y., P. Spring, S. Denev, and J. Sweetman (2007). Effect of a mannan oligosaccharide on the growth performance and immune status of rainbow trout (*Oncorhynchus mykiss*). Aquaculture International, 15, 2, 153–161.

Valtchev, I., Ts. Koynarski, L. Sotirov, Y. Nikolov, and P. Petkov (2015). Effect of aflatoxin B1 on Moulard duck’s natural immunity. Pakistan Veterinary Journal, 35, 67–70.

Yin, Y.L., Z.R. Tang, Z.H. Sun, Z.Q. Liu, T.J. Li, R.L. Huang, Z. Ruan, Z.Y. Deng, B. Gao, L.X. Chen, C.Y. Wu, and S.W. Kim (2008). Effect of galacto-mannan-oligosaccharides or chitosan supplementation on cytoimmunity and humoral immunity in early-weaned piglets, AJAS, 21, 5, 723–731.

Zhang, Sh., B. Liao, X. Li, L. Li, L. Ma, and X. Yan (2012). Effects of yeast cell walls on performance and immune responses of cyclosporine A-treated, immunosuppressed broiler chickens. Brittish Journal of Nutrition, 10, 858–866.

Zhang, Z.F., and I.H. Kim (2014). Effects of multistrain probiotics on growth performance, apparent ileal nutrient digestibility, blood characteristics, cecal microbial shedding, and excreta odor contents in broilers. Poultry Science, 93, 364–370.

